# A Bacterial Signaling Network Controls Antibiotic Resistance by Regulating Anaplerosis of 2-oxoglutarate

**DOI:** 10.1101/2020.10.22.351270

**Authors:** M. N. Hurst, C. J. Beebout, R. Mersfelder, A. Hollingsworth, K. R. Guckes, T. Bermudez, K. A. Floyd, S. A. Reasoner, D. Williams, M. Hadjifrangiskou

**Author notes:** Please address correspondence to Lead Contact: Maria Hadjifrangiskou, Division of Molecular Pathogenesis, Department of Pathology, Microbiology & Immunology, Vanderbilt University Medical Center, Nashville, TN, USA., Twitter: @BacterialTalk. Department of Biochemistry & Molecular Biology, Pennsylvania State University, Harrisburg, PA, USA. Department of Microbiology and Environmental Toxicology, University of California – Santa Cruz, Santa Cruz, CA, USA.

## Abstract

Antibiotic resistance has become a global threat. In addition to acquiring resistance via horizontal gene transfer, bacteria can evade killing by temporarily modifying their cell envelope to prevent antibiotic-bacterial interactions. A critical gap in knowledge is how bacteria balance the metabolic needs of altering the cell envelope with the constant need to generate energy. Cross-regulation between two signaling networks in *Escherichia coli* increases resistance to positively charged antibiotics. We show that increased resistance is supported by metabolic re-wiring controlled by the QseB transcription factor. QseB controls the increase in 2-oxoglutarate required for lipid A modification, by upregulating three anaplerotic pathways that feed acetyl Co-A, succinate and fumarate into the TCA cycle. Exogenous addition of 2-oxoglutarate restores antibiotic resistance in the *qseB* deletion mutant. Antibiotic resistant clinical isolates bear mutations within QseB-mediated anaplerotic pathways. These findings are significant, because they uncover a previously unknown mechanism of metabolic control of antibiotic resistance.

## Introduction

Enterobacteriaceae are common causative agents of disease in humans. Bacteria from this class account for urinary tract infections, bloodstream infections and pneumonias (Paterson 2006). To treat bacterial infections, antibiotics are indicated. However, antibiotic resistance in the Enterobacteriaceae is on the rise, with many strains harboring mobile genetic elements that code for extended spectrum β-lactamases (ESBL), which hydrolyze broad- and extended spectrum cephalosporins, monobactams, and penicillins (*1*). Posing an even greater challenge, many Enterobacteriaceae have now gained multi-drug resistance to other first line antibiotics. For example, oftentimes the genes that encode ESBLs are found on the same plasmid as genes that encode resistance to aminoglycosides and sulfonamides (*2*), while many other strains also possess the ability to resist quinolones. This creates a complex challenge to treat certain Gram-negative Enterobacteriaceae in the clinic. Grim statistics accompanying those of antibiotic resistance include high rates of antibiotic treatment failure, with longitudinal studies indicating that one in every ten antibiotic prescriptions fails even when the clinical laboratory’s antimicrobial susceptibility panel predicts susceptibility to a given drug regimen (*3-6*). The molecular underpinnings behind such treatment failures remain largely undefined. This work elucidates a previously uncharacterized mechanism in *Escherichia coli* that fuels resistance to positively charged antibiotics.

To treat infections caused by multi-drug resistant Enterobacteriaceae, alternative antibiotics are being used, including aminoglycosides and – to a lesser extent – polymyxins (*7*). For example, in urinary tract infections caused by ESBL- producing uropathogenic *E. coli* amikacin, a synthetic aminoglycoside, has shown good efficacy in treatment (*8*). Aminoglycosides and polymyxins are polycationic in nature and make contact with the bacterial cell envelope by binding to the lipopolysaccharide (LPS), phospholipids, and bacterial outer membrane proteins. This interaction leads increased permeability and penetration of the aminoglycoside or polymyxin into the periplasm. A mechanism used by bacteria to repel cationic antibiotics makes the bacterial cell envelope less negatively charged (*9*).

Lipopolysaccharide (LPS) comprises the outer leaflet of the outer membrane of *E. coli* and other Gram-negative bacteria. LPS is made up of a core lipid A anchor, core oligosaccharide and – in several strains – O antigen, all of which are synthesized at the inner leaflet of the inner membrane (**Figure 1A**). O-antigen and lipo-oligosaccharide are flipped to the periplasmic side of the inner membrane by the O-antigen flippase and the MsbA flippase respectively, where O-antigen is ligated onto the lipo-oligosaccharide by an enzyme called O-antigen ligase. The resulting LPS is transported to the outer membrane by the Lpt transport system (*10*). Lipids for new LPS are tightly regulated in synthesis of lipid A and arise from phospholipid biogenesis (*11*). Plasmid-borne determinants that confer resistance to polymyxins and other cationic polypeptides primarily code for enzymes that modify the charge on LPS, thereby reducing the electrostatic attraction of cationic molecules to the negatively charged bacterial cell envelope. However, most of the genes associated with LPS modification are also chromosomally encoded, enabling bacteria to transiently alter cell envelope charge without *a priori* acquisition of antibiotic resistance determinants. This ability contributes to the ability of bacteria to withstand antibiotic treatment (*12*).

**Figure 1:**
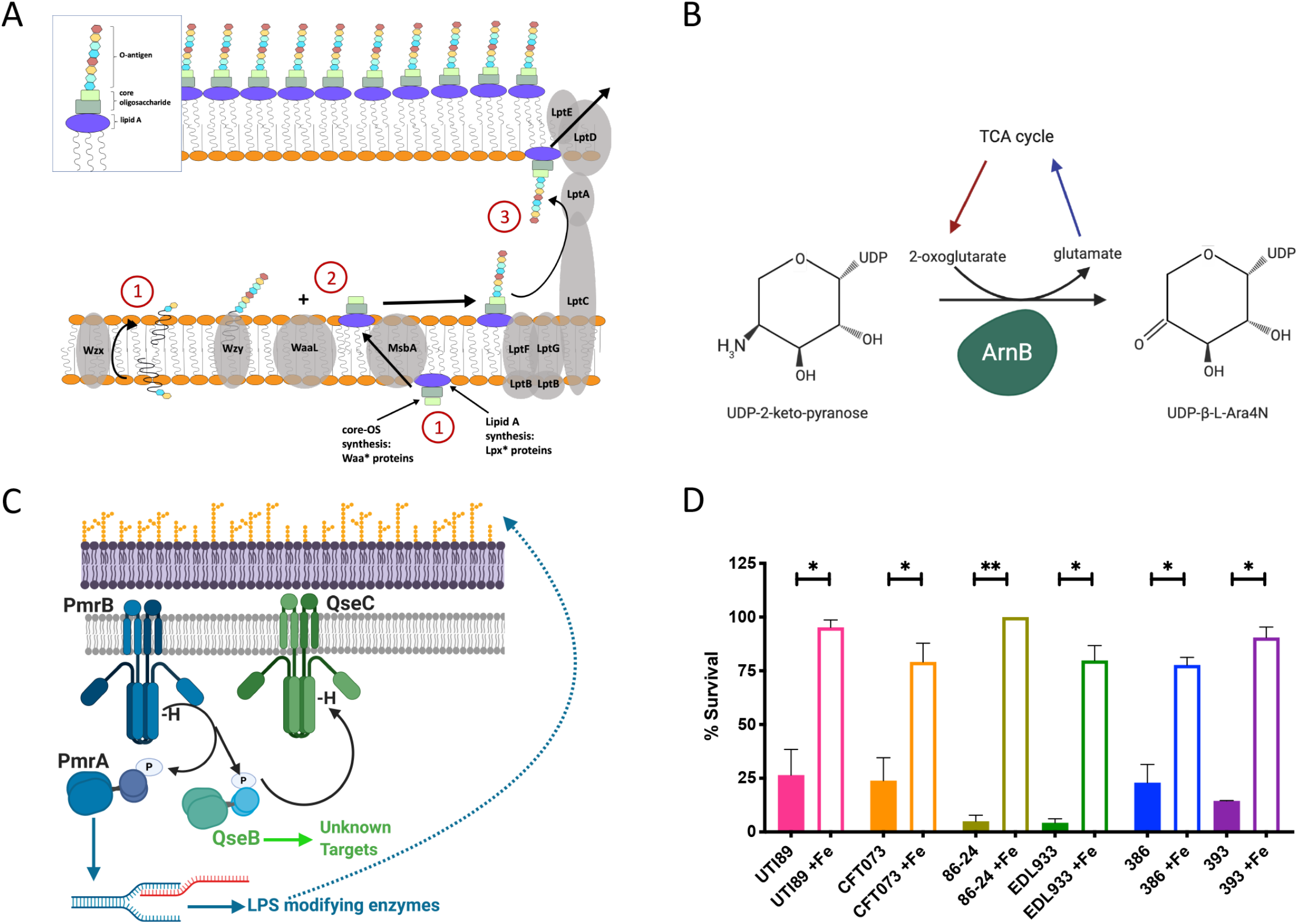
QseBC-PmrAB mediated control of polymyxin B resistance in *E. coli*. ((A) Cartoon depicts - in a simplified manner - the steps in lipopolysaccharide (LPS) biosynthesis in *Escherichia coli*. (B) Schematic shows the ArnB-mediated reaction that converts UDP-2-ketopyranose to UDP-b-L-Ara4N by consuming oxoglutarate and producing glutamate. (C) Cartoon depicts the mechanism of activation of the two-component systems PmrAB (blue) and QseC (green). PmrB is a membrane bound histidine kinase that is activated by ferric iron. Upon activation it auto-phosphorylates and then transfers the phosphoryl group onto its cognate response regulator PmrA and the non-cognate QseB. PmrA is a transcription factor, regulating the transcription of LPS modifying genes. QseB is also transcription factor, whose targets are unknown. (D) Graph depicts results of polymyxin B survival assays for each strain. Cells were allowed to reach mid logarithmic growth phase in the presence or absence of ferric iron and normalized. Cells were then exposed to polymyxin B at 2.5 μg/mL or to diluent alone (sterile water), for one hour. At this time cells were serially diluted and plated to determine colony forming units per mL. To determine percent survival, cells exposed to polymyxin were compared to the untreated controls (mean ± SEM, n = 3). To determine statistical significance, a Welch’s T-test was performed between the untreated strain and the same strain treated with ferric iron. (*), p < 0.05; (**), p < 0.01.

Altering the net charge of the envelope can be accomplished through different mechanisms, several of which target lipid A (*13*). The net charge of the membrane can be increased by dephosphorylation of lipid A, or addition of positively charged groups – such as phosphoethanolamine and amino-arabinose – directly to the lipid A group during synthesis (*14*). Addition of other groups, such as glycine, as well as lipid A acylation have also been observed to increase resistance to polymyxins and other cationic antimicrobial peptides. Finally, in addition to lipid A modifications, core-oligosaccharide and O-antigen components can also be modified through truncation, acylation, glycosylation, or addition of phosphoryl groups, colanic-, or sialic acids (*9*).

One fundamental question that remains unanswered is how *E. coli* and other LPS-containing microorganisms offset the metabolic burden associated with LPS modification. In *E. coli*, the majority of LPS modifications occur during new LPS biogenesis at the cytoplasmic leaflet of the inner membrane (*9*) and consume several products of central metabolism. This work focuses on consumption of 2-oxoglutarate by the chromosomally-encoded Arn* proteins, that add a 4-amino-4-deoxy-L-arabinose group onto nascent lipid A molecules. ArnA, ArnB, and ArnC convert undecaprenyl-glucuronic acid to undecaprenyl phosphate-*α*-4-amino-4-deoxy-L-arabinose, while ArnT catalyzes the transfer of the undecaprenyl phosphate-*α*-4-amino-deoxy-L-arabinose onto a nascent lipid A moiety. ArnB catalyzes a transamination reaction of undecaprenyl-4-keto-pyranose to undecaprenyl 4-amino-4-deoxy-L-arabinose by utilizing oxoglutarate and producing glutamate in the process (**Figure 1B**). How the cell fulfills the higher demand for oxoglutarate in this step remains unknown.

In previous work we determined that the PmrAB two-component system interacts – via phosphotransfer events – with QseB, another transcription factor that forms a two-component system with the QseC membrane-bound receptor (**Figure 1C** and (*12, 15, 16*)). Specifically, we discovered that activation of the PmrB receptor by one of its ligands – ferric iron – leads to phosphorylation of both the cognate PmrA and the non-cognate QseB and that both phosphorylation events are necessary for *E. coli* to mount resistance to polymyxin B (*12*). Deletion of the *pmrB* receptor abolishes the ability of *E. coli* to survive polymyxin intoxication; deletion of either *pmrA* or *qseB* leads to a two- to ten-fold reduction in survival, with the double deletion mutant Δ*pmrAΔqseB* phenocopying the *pmrB* receptor mutant (*12*). While PmrA regulates the expression of LPS-modifying enzymes in both *Salmonella* and *E. coli* (*17, 18*), the precise role of QseB in mediating resistance to polymyxin B in *E. coli* is unknown. Intriguingly, studies have recently reported the presence of an additional *qseBC* locus within an *mcr*-containing plasmid in a colistin-resistant isolate (*19*) further suggesting a role for the QseBC two-component system in LPS modification.

In this work, we demonstrate that the QseB transcription factor mediates resistance to a range of positively charged antibiotics – gentamicin, amikacin and polymyxin B – by controlling the anaplerosis of oxoglutarate. QseB controls the metabolism of glutamate via three routes that feed succinate, fumarate and Co-enzyme A into the tricarboxylic acid (TCA) cycle. Fueling the TCA cycle at points past the succinyl-CoA step serves to mitigate the loss of oxoglutarate to LPS biogenesis, allowing the TCA cycle to continue running during production of modified LPS. Accordingly, deletion of *qseB* leads to increased glutamate levels during antibiotic intoxication, a phenomenon that is abolished in the *qseB* complementation strain or upon exogenous addition of oxoglutarate. Deletion of QseB-regulated genes involved in oxoglutarate production or glutamate utilization also phenocopy the *qseB* deletion phenotype. Deletion of a bacterial oxoglutarate sensor also attenuates antibiotic resistance. Finally, analyses of clinical isolates with stable polymyxin B resistant subpopulations identified in patients with urinary tract infections sustain polymorphisms in succinate and oxoglutarate metabolism enzymes. We propose a new role for QseB in balancing the metabolic requirements associated with LPS modification. These findings are significant, because they uncover a previously unknown mechanism of metabolic control of antibiotic resistance.

## Results

### QseBC-PmrAB signaling is active in prevalent *E. coli* clades

In uropathogenic *E. coli* (UPEC), resistance to polymyxin B is mediated through the concerted action of two interacting two-component systems <u>**P**</u>oly<u>**m**</u>yxin <u>**r**</u>esistance (Pmr)AB and **Q**uorum **Se**nsing (Qse)BC (*12, 15, 16, 20*). In *Salmonella* spp., PmrAB orchestrates the activation of genes involved in lipid A modification in response to elevated concentrations of ferric iron, which are sensed by the PmrB sensor histidine kinase (*21, 22*). Activation of PmrB via auto-phosphorylation results in phosphotransfer to the cognate response regulator PmrA, which transcriptionally regulates several genes involved in LPS modification (*13, 22, 23*). In UPEC, activation of PmrB by increased levels of ferric iron leads to auto-phosphorylation and subsequent phosphotransfer to two response regulators: the cognate PmrA and the non-cognate QseB (*12, 16, 20*). Deletion of either *qseB* or *pmrA* leads to a 2- to 10-fold reduction in the ability of *E. coli* to resist killing by Polymyxin B (PMB), while deletion of both response regulators phenocopies the *pmrB* deletion which is nearly 100% sensitive to PMB (*12*). The QseC sensor histidine kinase – the cognate partner of QseB – is critical for system de-activation (*20*).

We first asked whether QseBC is involved in mediating polymyxin B resistance in diverse *E. coli* strains. There are five prevalent phylogenetic clades that host the majority of sequenced *E. coli* isolates, A, E, B2, D, B1. Using a representative panel of *E. coli* strains that lack plasmid-borne polymyxin B resistance determinants and are sensitive to polymyxin, based on clinical testing (**Figure 1D**), we asked if pre-treatment with the PmrB ligand ferric iron would induce activation of the *qseBC* promoter, which is under the control of QseB (*20, 24-27*). Transcriptional profiling by reverse transcription-followed by quantitative qPCR revealed a robust activation surge (*28*) of the *qseBC* promoter in all strains tested from clades A, E, B2, D and B1 (**Figure S1**). Evaluation of strain survival in 2.5X the Minimum Inhibitory Concentration (MIC) of polymyxin B revealed that all tested strains exhibited 77 – 100% survival compared to untreated controls which exhibited 4 – 22% survival. (**Figure 1D**). Deletion of *qseB* or *qseBC* in well-characterized enterohemorragic strains 86-24 and 87-14 led to a significant reduction in polymyxin B resistance compared to the wild-type parent (**Figure S2**), which was rescued upon extra-chromosomal complementation with a wild-type copy of *qseB*. Notably, strain Sakai harbors a truncated, non-functional copy of QseC, a phenomenon which would suggest that in this strain background, PmrB-QseB interactions are unrestrained as previously seen for a uropathogenic *E. coli* strain deleted for *qseC* (*15*). In UPEC strains, unrestrained interaction of PmrB and QseB leads to increased resistance to polymyxin B in a signal-independent manner (*12*). Indeed, strain Sakai exhibits intrinsic resistance to polymyxin B (**Figure S2)**, consistent with the model that absence of functional QseC leads to uncontrolled PmrB-to-QseB phosphotransfer and subsequent intrinsic resistance to polymyxins. Supporting this notion, deletion of *qseB* or the entire *qseBC* locus in this strain phenocopies the *qseB* deletion in the other EHEC isolates (**Figure S2**). Combined these results indicate that transient resistance to polymyxin B in diverse *E. coli* clades is mediated at least in part by PmrB – QseB interactions. To further probe the the mechanism by which QseB mediates antibiotic resistance we used the prototypical pathogenic strain UTI89.

### QseB mediates resistance to positively charged antibiotics

If QseB plays a critical role in supporting modification of the cell envelope to resist polymyxin B intoxication, one would expect that QseB activation would lead to resistance to a broad range of positively charged antibiotics. To test this hypothesis, isogenic strains lacking QseB, or carrying QseB in the native locus or extra-chromosomally, were tested for their ability to resist gentamicin or amikacin, two aminoglycoside antibiotics that are positively charged. Nitrofurantoin, which is neutral, along with polymyxin B were used as controls. Strains were tested for their ability to survive a concentration of antibiotic at up to 5 times the established minimum inhibitory concentration – in conditions without or with preconditioning with ferric iron. These studies indicated that the strains harboring QseB (wild-type strain (**Figure 2A, 2D, 2G**) and the Δ*qseB*/pQseB complemented strain (**Figure 2C, F, I**)) exhibited 85-95% survival in the presence of charged antibiotics when pre-conditioned with ferric iron (**purple lines, Figure 2)**. However, the strain lacking qseB (Δ*qseB*, **Figure 2B, E, H)** exhibited a marked decline in survival regardless of the presence of ferric iron. Similarly, the uncharged antibiotic nitrofurantoin led to effective bacterial killing of all genetic backgrounds and bacterial survival was not affected by the presence of QseB (**Figure 2J, K, L**). Combined, these data demonstrate that the QseB transcription factor mediates resistance to positively charged antibiotics.

**Figure 2:**
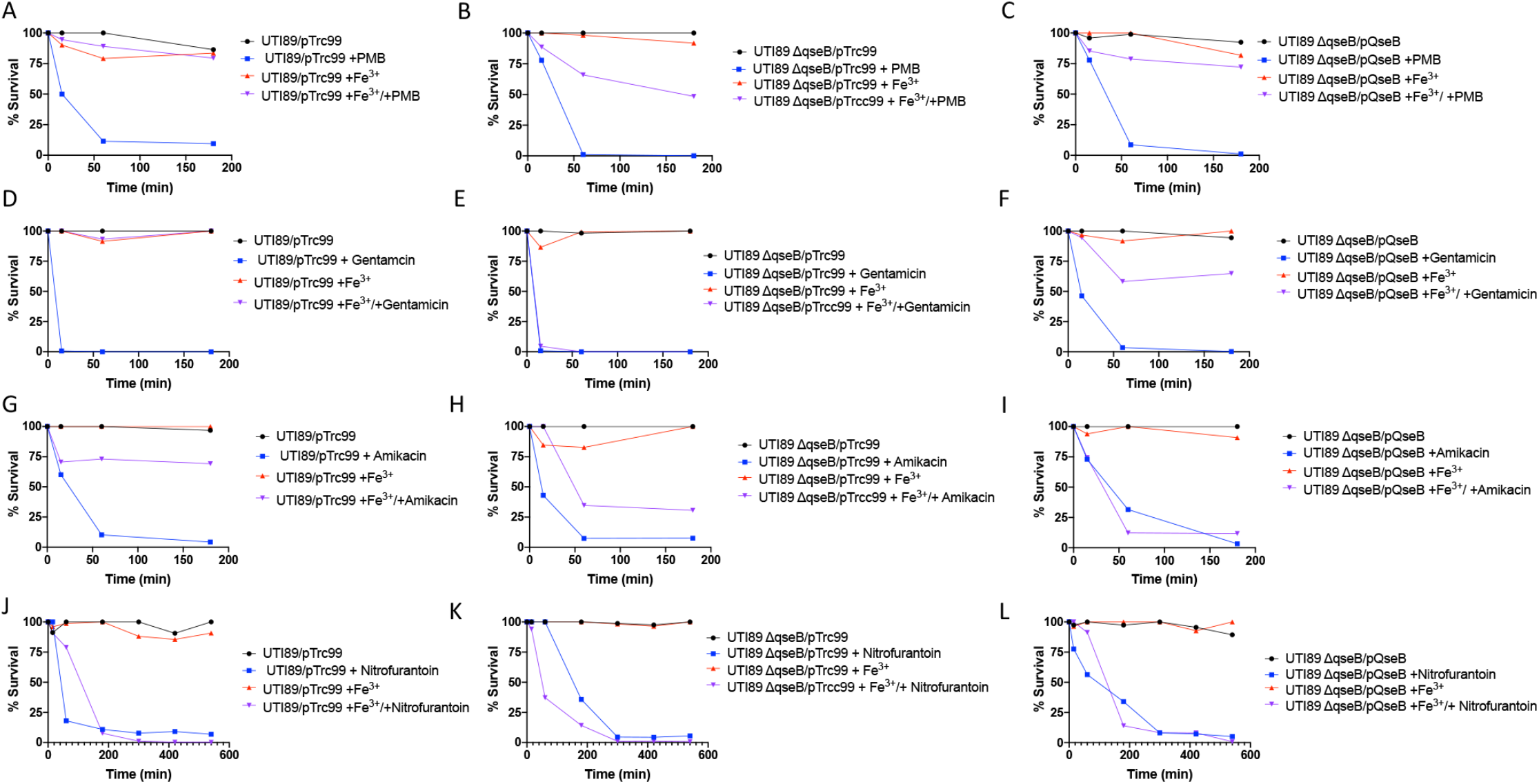
QseB confers resistance to positively charged antibiotics. Graphs depict survival across time during antibiotic challenge of isogenic strains lacking or harboring QseB. In each experiment, strains were challenged with: no additives (Black lines, control); 100 μM ferric iron (Red lines; PmrB activation stimulus); antibiotic of choice (Blue lines), or; both 100 μM ferric iron and an antibiotic of choice (Purple lines). (A-C) Polymyxin survival assays with UTI89/pTrc99 (A); UTI89Δ*qseB*/pTrc99 (B), or UTI89Δ*qseB*/pQseB. (D-F) Gentamicin survival assays with UTI89/pTrc99 (D); UTI89Δ*qseB*/pTrc99 (E), or UTI89Δ*qseB*/pQseB (F). (G-I) Amikacin survival assays with UTI89/pTrc99 (G); UTI89Δ*qseB*/pTrc99 (H), or UTI89Δ*qseB*/pQseB (I) (J-L) Nitrofurantoin survival assays with UTI89/pTrc99 (J); UTI89Δ*qseB*/pTrc99 (K), or UTI89Δ*qseB*/pQseB (L). All graphs show a representative of three trials (n = 3).

### Temporal Tracking of Gene Activation Under PmrB-Activating Conditions Reveals an Anaplerotic Circuit Under QseB Control

Previous studies in *Salmonella* and *E. coli* elucidated PmrA targets that are responsible for modification of the LPS (*13, 22, 23, 29*). Previous studies in *E. coli* revealed that deletion of *qseC* confers a significant shift in the expression of genes involved in central metabolism, including – among others – genes involved in glutamate metabolism and the tricarboxylic acid cycle (*26*). To decipher the regulon of QseB under conditions of antibiotic stress, steady-state transcript abundance across the activation surge were tracked over time via RNA sequencing (RNAseq). In parallel, promoters bound by QseB were identified using Myc-His-tagged QseB and crosslinking, followed by immunoprecipitation and analysis of bound DNA sequences on microarray chips (chIP-on-chip).

For the RNAseq experiments, the wild-type strain UTI89 and isogenic mutants lacking *qseB* (UTI89Δ*qseB*, or *pmrA* and *qseB* (UTI89Δ*pmrAΔqseB*) were grown under PmrB-activating conditions (100μM Ferric iron, Fe^3+^) and samples were obtained for RNA sequencing immediately prior to (T=0), as well as 15 (T=15) and 60 (T=60) minutes post addition of ferric iron to the growth medium (**Figure 3A)**. Output RNA sequencing data from three biological repeats per strain per timepoint were analyzed using Rockhopper software (*30, 31*). Differential gene expression matrices within each strain were calculated to compare T = 0 to T =60 and T =15 minutes (**Figure 3B-C and Supplementary File 1**). An additional comparison of T=15 and T=60 was also made (**Supplementary File 1**). Transcripts with a q value lower than 0.05 were considered significant (**Supplementary File 1**). The RNAseq profile of the wild-type strain, UTI89, revealed a total of 829 transcripts with significantly altered abundance across time, in response to the addition of ferric iron (**Supplementary File 1, worksheet 1**). Of these, 49 belonged to small non-coding RNAs, while 148 were hypothetical proteins (**Supplementary File 1, worksheet 1;** highlighted in light- and dark gray respectively). Of the remaining 632 transcripts, 81 belonged to tRNAs (n=50) and transcripts coding for tRNA modification- or translation-associated proteins (n=31) (**Supplementary File 1, worksheet 1;** highlighted in light pink). The non-coding RNA transcripts, as well as hypothetical and tRNA-/translation associated proteins were excluded from the heatmaps, along with 6 plasmid-associated transcripts unique to UTI89. The remaining 545 transcripts are depicted in heatmaps, according to the pathways they belong to (**Figure 3B-C, Supplementary Figure S3**).

**Figure 3:**
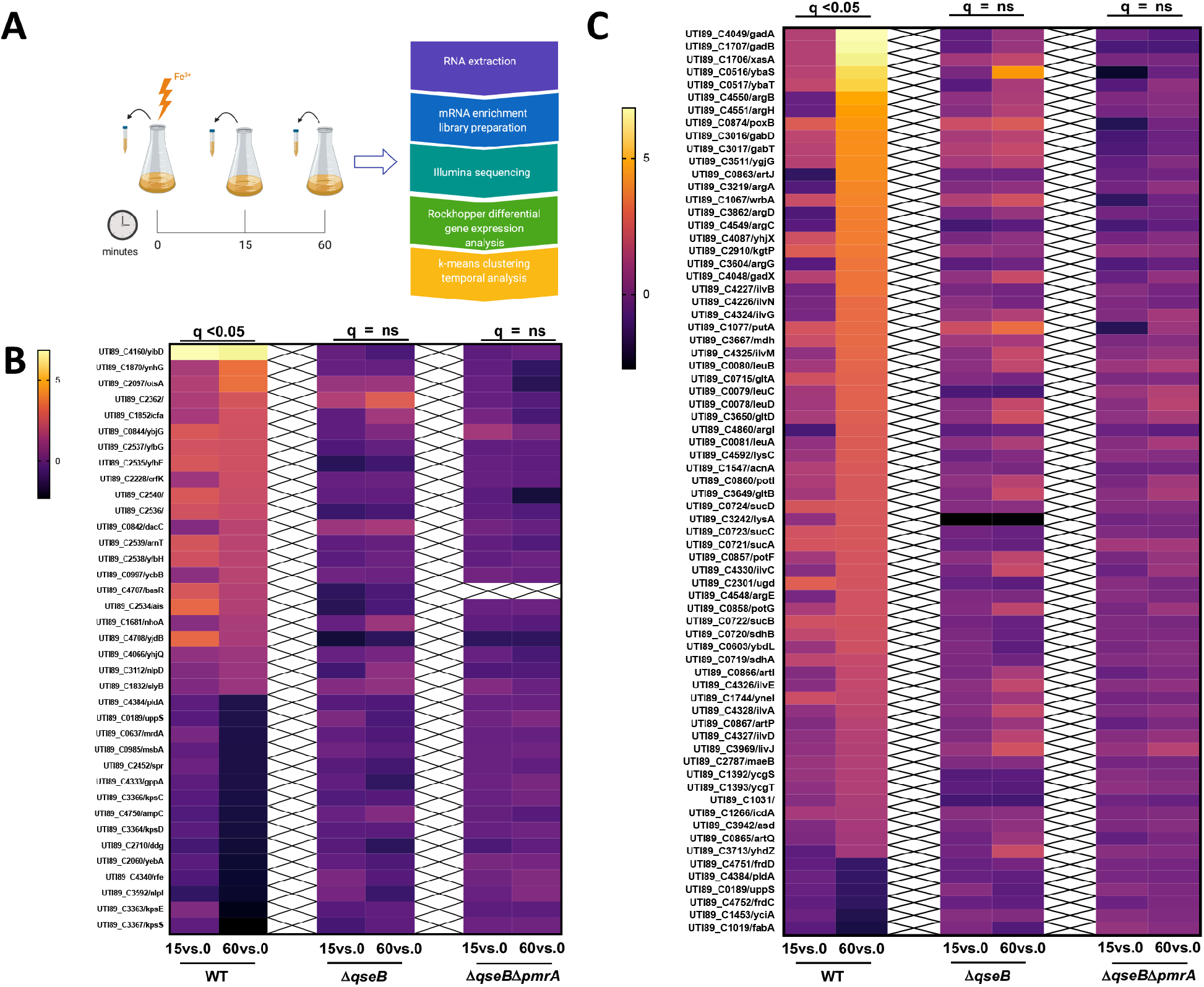
RNAseq profiling across time reveals a metabolic circuit controlled by QseB and PmrA. (A) Schematic showing the pipeline for sample collection and data processing. (B-C) Heatmaps indicate log2 relative fold change of UTI89 WT, Δ*qseB*, and Δ*qseBΔpmrA* for genes involved in cell envelope modification (B) or metabolism (C) after stimulation with ferric iron at 15- and 60-minutes post stimulation. These genes were significantly (q<0.05) changed at 60 minutes compared to pre-stimulation (T=0) in wild-type UTI89. (C) A heatmap showing log2 relative fold change of UTI89 WT, Δ*qseB*, and Δ*qseBΔpmrA* for genes in metabolic pathways after stimulation with ferric iron at 15- and 60-minutes post stimulation. These genes were significantly (q<0.05) changed at 60 minutes compared to pre-stimulation (T=0) in wild-type UTI89, but not in the mutant strains.

Although LPS modification genes surged over time, following stimulation with ferric iron (**Figure 3B-C**), the most highly upregulated genes in response to ferric iron stimulation were genes that belonged to glutamate metabolism and the TCA cycle (**Figure 3C**). The same clusters had no significant surge in the mutants lacking QseB (**Figure 3C-D, Supplementary File 1**). These data indicate that QseB and PmrA are functionally redundant in mediating transcription of LPS modification genes in pathogenic *E. coli*. Furthermore, these data indicate a role for QseB in mediating glutamate metabolism.

### ArnA/B and enzymes involved in glutamate metabolism, pantothenate and coenzyme A synthesis are under the control of QseB

To determine promoters bound by QseB, the UTI89□Δ*qseB* strain that harbors a construct expressing Myc-His-tagged QseB under an arabinose-inducible promoter (*12, 15, 27*) was subjected to chIP-on-chip analyses, using UTI89-specific Affymetrix chips (*26*). An isogenic strain harboring the pBAD-MycHis A vector was used as a negative control. Cultures were grown in laboratory media in the presence of 0.02□M arabinose to ensure constant expression of QseB, at concentrations similar to those we previously published as sufficient for QseBC complementation (*15*). Pull-downs using an anti-Myc antibody were performed on six separate reactions, three for the experimental and three for the control strain. Analyses of the pull-down DNA revealed a total of 169 unique promoters bound by QseB and absent in the negative control (**Figure S4** and **Supplementary File 1**). Sixty-three out of the 169 promoters were also reported in the microarray analyses of the *qseC* mutant in previous studies (*26*). Among the promoters identified was the promoter of *qseBC* – consistent with QseB’s ability to regulate its own transcription (*15, 24, 26*). Another set of targets included the promoters of *arnB* (*yfbE*), *argF, asnB*, the *ilv* cluster, *panBCD* and *sucD* and *asnB* (**Figure S4)**, as well as *glnK, glnS* and *aspS* that could indirectly be responsible for the transcriptional effects observed in the RNAseq analyses. (**Figure 3C** and **Supplementary File 1**). The genes directed by these identified promoters are involved in nitrogen assimilation, glutamate metabolism, panthothenate and coenzyme A synthesis. As mentioned above, the *arnB* gene, encodes the enzyme that catalyzes transamination of undecaprenyl-4-keto-pyranose to undecaprenyl 4-amino-4-deoxy-L-arabinose (**Figure 1B**). This ArnB-mediated reaction consumes oxoglutarate and produces glutamate. Intriguingly, the QseB-bound promoters identified in the chIP-on-chip analyses (**Figure S4, Supplementary File 1**), along with the RNAseq data (**Figure 3C**) are connected through their involvement in glutamate metabolism (**Figure 4**).

**Figure 4:**
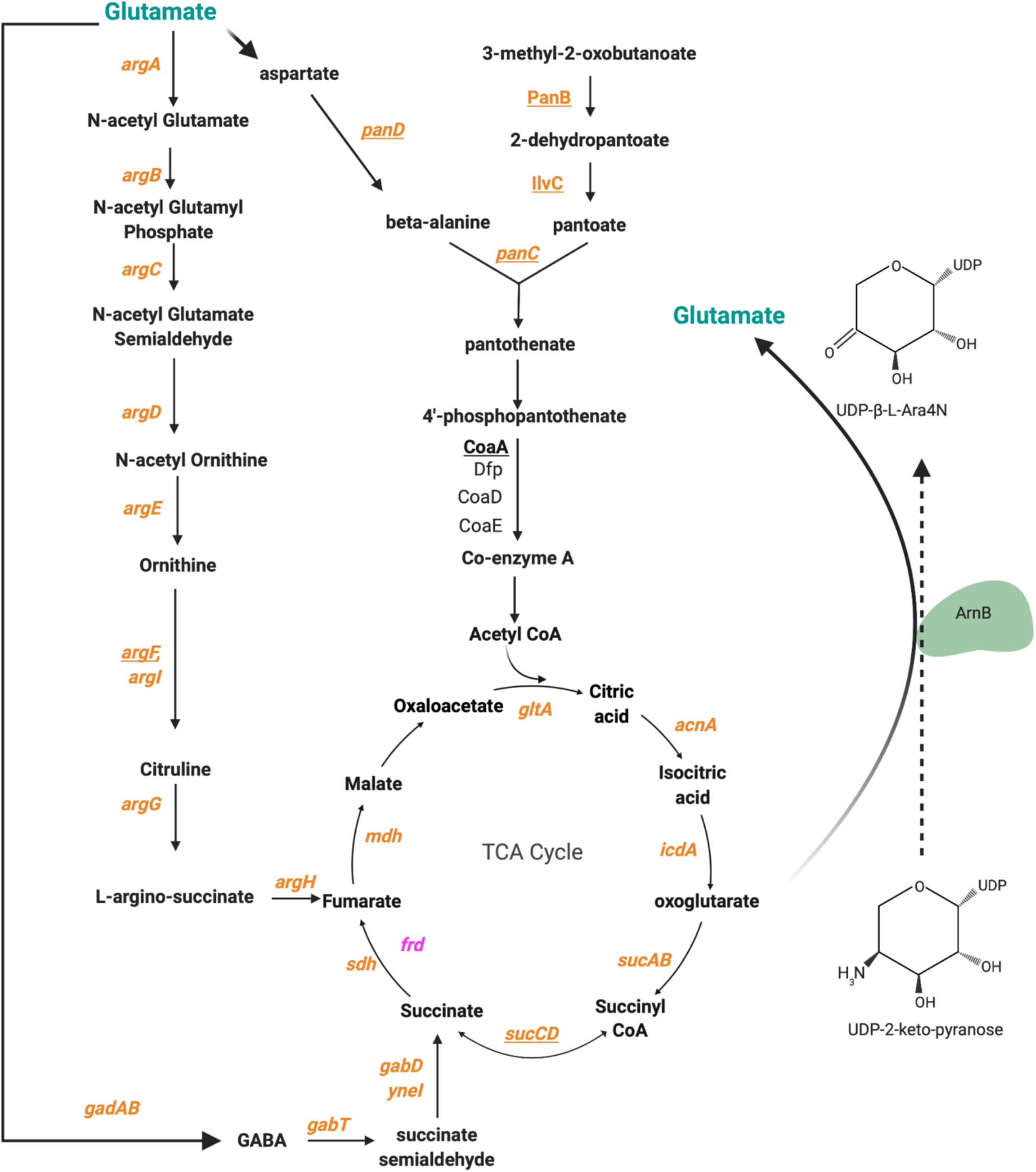
The anaplerotic circuit controlled by QseB. Schematic outlining the metabolic pathways that QseB interacts with. Genes in orange are upregulated at 60 minutes post-stimulation with ferric iron as indicated by RNA sequencing data. Genes in pink are downregulated at 60 minutes post-stimulation with ferric iron. Genes underlined are direct targets of QseB as indicated by ChIP-on-chip data.

### Glutamate, aspartate and Co-enzyme A concentrations change in the absence of QseB

To determine whether QseB plays a role in glutamate-oxoglutarate homeostasis during the ArnB-mediated LPS modification step, the levels of metabolites along the above identified pathway (**Figure 4**) were measured. Oxoglutarate is difficult to quantify, due to rapid degradation (*32*), thus oxoglutarate measurements were not taken. For metabolomics measurements, samples were obtained at the same time intervals shown for the RNA sequencing analyses (**Figure 3A**). Glutamate (**Figure 5A-C**), aspartate (**Figure 5D-F**), coenzyme A (**Figure 5G-I**) measurements were taken in the wild-type parent and the isogenic Δ*qseB* and complemented strains, in the presence of ferric iron alone (**Figure 5**, red lines) polymyxin B (PMB) alone (**Figure 5**, blue lines), ferric iron + PMB (**Figure 5**, pink lines) or no additives (**Figure 5**, black lines). In each panel, concentrations were normalized to the unstimulated/unchallenged control (black lines). In the wild-type background, addition of polymyxin B and ferric iron resulted in a decrease in glutamate levels compared to cells exposed to ferric iron alone or no additives (**Figure 5A**). Conversely, in a *qseB* deletion mutant, glutamate levels remain relatively unchanged following addition of polymyxin B and ferric iron compared to ferric iron alone or with no additives (**Figure 5B**). Glutamate levels decrease again shortly after addition of ferric iron and polymyxin B in a *qseB* deletion mutant complemented with a wild-type copy of *qseB* (Δ*qseB*/pQseB, **Figure 5C**). These data indicate that glutamate levels change in response to polymyxin B addition in a manner that depends upon QseB. This could suggest that rapid glutamate utilization occurs, following excess production during the conversion of undecaprenyl-4-keto-pyranose to undecaprenyl 4-amino-4-dexoy-L-arabinose.

**Figure 5:**
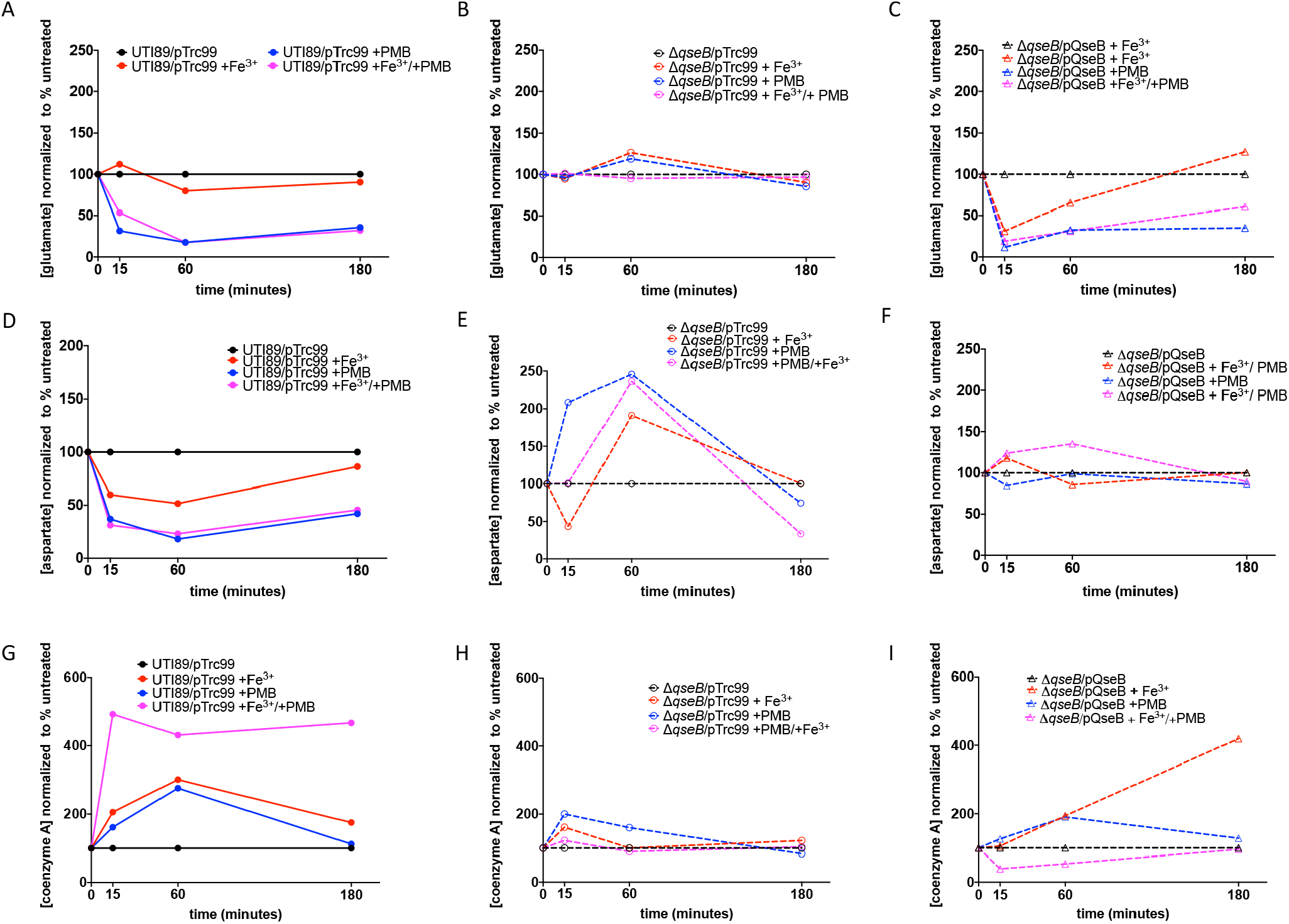
Metabolomic changes in response to LPS modification and QseB control. Graphs indicate metabolite abundance for glutamate, aspartate and co-enzyme A over time in wild-type pathogenic E. coli and isogenic mutants under different stimulation conditions. Measurements are normalized to a sample in which no additives or conditions were changed (black lines). Pink lines show measurements from samples in which ferric iron and polymyxin were added. Blue lines show measurements from samples in which only polymyxin was added. Red lines show measurements in which only ferric iron was added. **(A-C)** Glutamate was measured across time in wild-type UTI89 (A), UTI89Δ*qseB*/pTrc99 (B) and UTI89Δ*qseB/pQseB* (C). (**D-F**) Aspartate was measured in wild-type UTI89 (D), UTI89Δ*qseB*/pTrc99 and (E) UTI89Δ*qseB/pQseB* (F). (G-I) Aspartate was measured in wild-type UTI89 (G), UTI89Δ*qseB*/pTrc99 (H), and UTI89Δ*qseB/pQseB* (I). Graphs are representative of at least three biological repeats.

Because glutamate can be converted to aspartate via the action of AspC (**Figure 5**), we also measured aspartate levels in the same samples (**Figure 5D-F**). Similar to glutamate, aspartate levels dropped in wild-type samples treated with polymyxin B and ferric iron compared to a sample with no additives (**Figure 5D**). In a *qseB* deletion mutant aspartate levels rise after addition of polymyxin B and ferric iron compared to a sample with no additives. In a *qseB* deletion complemented with *qseB* aspartate levels remain relatively unchanged compared to a sample with no additives (**Figure 5F**). Aspartate feeds into coenzyme A production through the pantothenate synthesis pathway (**Figure 4**), of which several gene products involved are direct targets of QseB (**Figure 4**). Following addition of polymyxin B and ferric iron, coenzyme levels rapidly rise (**Figure 5G**) in a wild-type sample, compared to a sample with no additives. However, in the *qseB* deletion mutant coenzyme A levels remain relatively unchanged or rise slightly (**Figure 5H)**. Interestingly in this case, complementation with pQseB did not complement the phenotype (**Figure 5I**).

### Oxoglutarate rescues the antibiotic susceptibility of the *qseB d*eletion mutant

The combined metabolomics, along with the RNAseq data suggest that QseB controls metabolic reactions that shunt the glutamate produced from the modification of LPS towards production of co-enzyme A, which could then re-enter the TCA (**Figure 4**), replenishing oxoglutarate levels. Moreover, if the RNAseq/chIP-on-chip observations truly map a metabolic regulon under QseB control in response to LPS modification, it appears that pantothenate production is also part of this process (**Figure 4**). To determine whether indeed these suggested anaplerotic pathways are critical in mounting a response to positively charged antibiotics, two mutants were created with deletions in *panD* and *icdA* respectively (**Figure 4**). PanD codes for an aspartate decarboxylase that converts aspartate into beta-alanine, which then feeds into the pantothenate pathway eventually resulting in coenzyme A production (**Figure 4**). The *pan* gene operon is under the direct control of QseB (**Figure S4 and Supplementary File 1**). We reasoned that if the identified QseB regulon is active during LPS modification, then deletion of *panD*, which is centrally placed in the identified pathway (**Figure 4**) should impair resistance to polymyxin B. To determine whether the pathway feeds into the TCA cycle, we also created an *icdA* deletion mutant, disrupting the conversion of isocitrate to oxoglutarate (**Figure 4**), limiting oxoglutarate amounts for LPS modification. Obtained mutants were tested in polymyxin B survival assays alongside the wild-type parental strain and the isogenic *qseB* deletion mutant, as well as the *qseB* deletion mutant complemented with *qseB*. Strains were tested for their ability to survive a concentration of PMB at five times the established minimum inhibitory concentration – as determined in **Figure 2A-C** – in conditions without or with preconditioning with ferric iron. While the wild-type and the Δ*qseB*/pQseB complemented strains exhibited 85-95% survival in 5X the PMB MIC when pre-conditioned with ferric iron (**Figure 6A**), the *qseB, panD* and *icdA* deletion mutants reproducibly exhibited a 50% reduction in survival.

**Figure 6:**
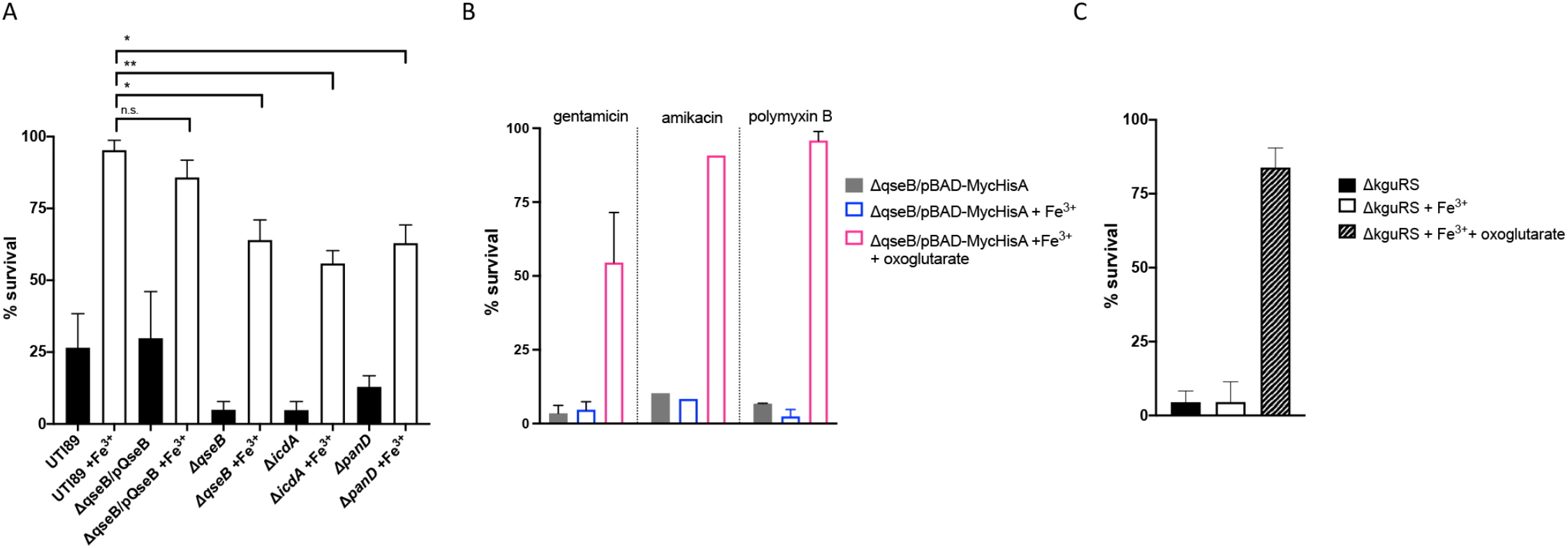
Oxoglutarate restores antibiotic resistance in the absence of QseB or an oxoglutarate sensor. **(A)** Graph depicts polymyxin B survival assays for wild-type *E. coli* and isogenic mutants deleted for *qseB, icdA*, or *panD*. Cells were allowed to reach mid logarithmic growth phase in the presence or absence of ferric iron and normalized. Cells were then either exposed to polymyxin at 2.5 μg/mL or without addition for one hour. To determine percent survival, cells exposed to polymyxin were compared to those that were not (mean ± SEM, n = 3). To determine statistical significance, a Welch’s T-test was performed between strains treated with ferric iron and UTI89 wild-type treated with ferric iron. (*), p < 0.05; (**), p < 0.01; (***), p <0.001. N.S. denotes a comparison that did not result in statistical significance. **(B)** Graph depicts gentamicin, amikacin and polymyxin B survival assays for *E. coli* isogenic mutants deleted for *qseB* and carrying the pBAD-MycHisA plasmid. Cells were allowed to reach mid logarithmic growth phase in the presence or absence of ferric iron and normalized. Cells were then exposed to gentamicin at 3.125 μg/mL, amikacin at 10 μg/mL or polymyxin B at 2.5 μg/mL or without addition for one hour. Cells were also exposed to ferric iron (100 μM) and/or oxoglutarate (5 mM). To determine percent survival, cells exposed to antibiotic were compared to those that were not (mean ± SEM, n =3 for gentamicin and polymyxin B assay, n = 1 for amikacin assay). (**C)** Graph depicts polymyxin B survival assay for *E. coli* isogenic mutants of *kguRS* an oxoglutarate sensing two-component system. Cells were allowed to reach mid logarithmic growth phase in the presence or absence of ferric iron and normalized. Cells were then exposed to polymyxin at 2.5 μg/mL or without addition for one hour. Cells were also exposed to ferric iron (100 μM) and/or oxoglutarate (5 mM). Percent survival was determined as described above and in the materials and methods.

If QseB controls the oxoglutarate “budget”, then susceptibility to charged antibiotics in the *qseB* deletion strain may be due to limitation in the production of oxoglutarate. To test this hypothesis, we asked whether the susceptibility of a *qseB* deletion mutant to positively charged antibiotics could be rescued via the addition of exogenous oxoglutarate. Remarkably, addition of oxoglutarate to ferric iron/polymyxin B-treated samples of Δ*qseB* restored survival in 5X the antibiotic MIC (**Figure 6B**). These data indicate that oxoglutarate anaplerosis controlled by QseB is necessary for mounting resistance to positively charged antibiotics.

Previous studies identified a two-component system, KguRS, prevalent in uropathogenic *E. coli* isolates that senses oxoglutarate (*33*). Using this system as an orthologous tool to validate the requirement for oxoglutarate, we asked whether a mutant lacking *kguRS* would have defects in modulating antibiotic resistance. We created a mutant lacking *kguRS* in wild-type UPEC and tested its susceptibility to PMB. The resulting mutant UTI89Δ*kguRS* demonstrated remarkable susceptibility to PMB (**Figure 6C**), a phenomenon that was rescued by exogenous addition of oxoglutarate (**Figure 6C**).

Together, these results indicate that oxoglutarate anaplerosis, as controlled by the QseB transcription factor is critical for transient resistance to positively charged antibiotics. Furthermore, these observations uncover an intrinsic mechanism by which *E. coli* can survive an antibiotic treatment regimen, if a subpopulation of bacteria has the PmrB-QseB-PmrA network activated.

### Clinical isolates with polymyxin hetero-resistance harbor mutations in QseB-regulated anaplerotic pathways

In clinical practice, up to 10% of prescribed regimens fail (*4-6, 34*), despite antibiogram outputs from the clinical laboratory that indicate bacterial susceptibility to the prescribed antibiotic regimen. Our observations suggest an intrinsic mechanism that may contribute – at least in part – to treatment failure in the case of pathogenic *E. coli* infections. To begin to address this hypothesis, we first evaluated the stochastic expression of the *qseB* locus in a bacterial community, without the presence of the ferric iron stimulus. Utilizing a previously created promoter construct fused to GFP, we assessed the activity of the promoter within bacterial biofilm communities, which are formed at different niches during infection (*35*). Microscopy revealed the presence of cells in which Pqse::GFP was active (**Figure 7A**), indicative of the presence of subpopulations within a bacterial community that have the potential to resist treatment with positively charged antibiotics.

**Figure 7:**
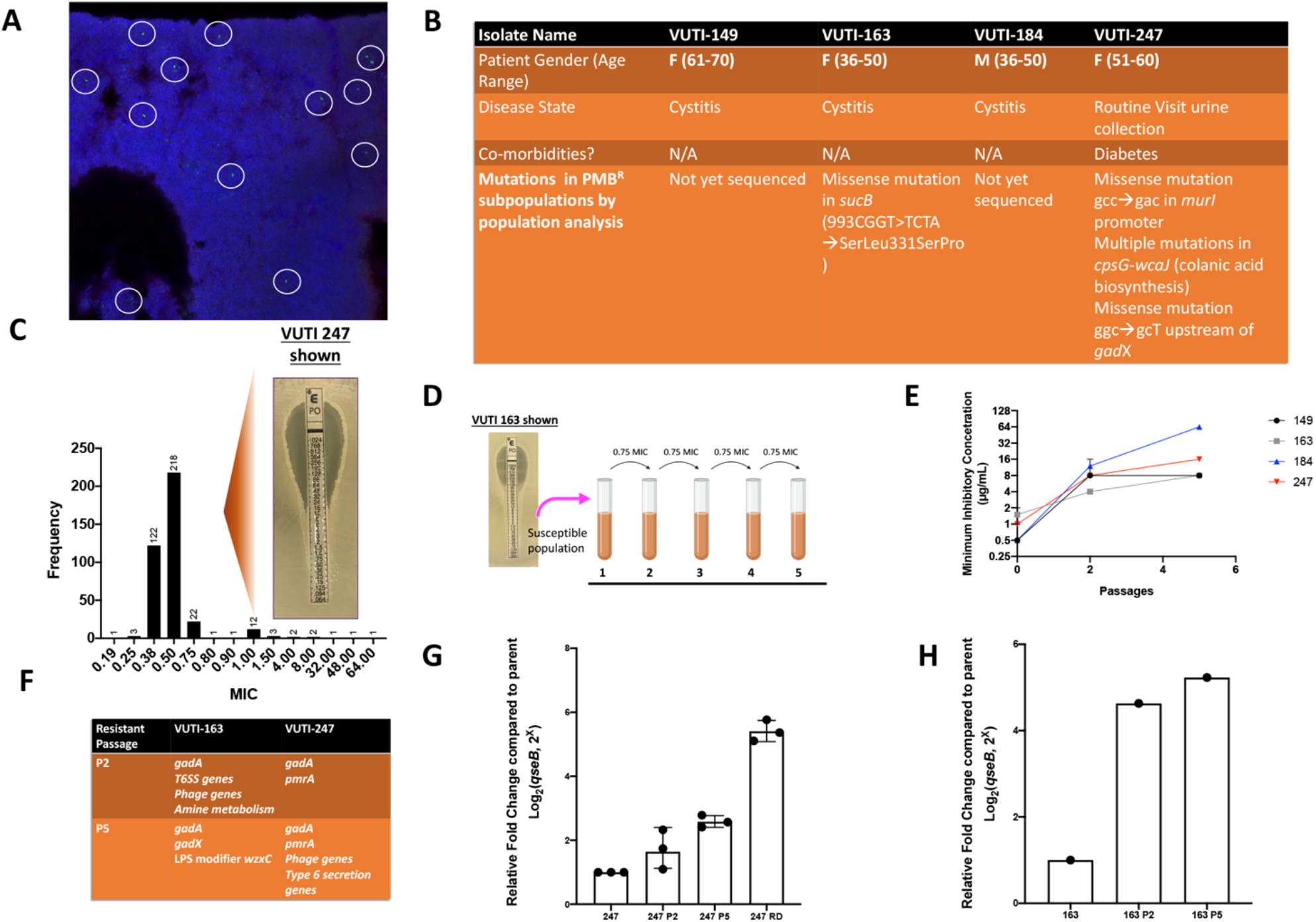
SNPs associated with polymyxin B-resistant subpopulations in clinical *E. coli* isolates. **(A)** Confocal Laser Scanning Microscopy (CLSM) images of *E. coli* biofilms formed by bacteria harboring the previously constructed P*qse*::GFP reporter. The P*qse*::GFP reporter is expressed in a small number of cells (marked with circles). Blue coloration is Topro-3 DNA stain. Images depicted are representative of at least 5 biological replicates. (**B)** Table depicts demographics associated with the strain source patients. Whole genome sequencing of VUTI 247 and VUTI163 reveals mutations compared to the susceptible subpopulation (see also supplementary file**). (C)** Graph shows the distribution of polymyxin B MICs for 300 *E. coli* urinary tract isolates collected at Vanderbilt University Medical Center Clinical laboratory under IRB#151465. MICs were determined using polymyxin E-test strips. Inset image depicts VUTI 247 exhibiting two populations with differing MIC’s: a major subpopulation with a susceptible MIC and a minor, yet stable, subpopulation with a resistant MIC. 2). **(D)** Schematic depicts the set-up for an *in vitro* evolution experiment. The susceptible population from each tested isolate demonstrating two populations (VUTI 149, VUTI 163, VUTI 184 and VUTI 247), was passaged 5 times, over a 5-day period, at 0.75X its polymyxin B MIC. **(E)** The MIC prior to passage, the second passage, and the fifth passage was determined using broth microdilution. Graph depicts the minimum inhibitory concentration of a strain over each passage (mean ± SEM, n = 3). **(F)** Table contains the mutations that were found after passages 2 and 5 of VUTI 163 and VUTI 247 underwent whole genome sequencing and were compared to the parent strains (See also supplementary file 2). **(G)** VUTI247, passage 2 and 5 and the resistant daughter from the resistant subpopulation were tested for activity of the *qse* operon. Cells were collected from a culture grown overnight allowed to reach stationary phase. RNA was extracted and reverse –transcribed. cDNA resulting from the reaction was subjected to qPCR with a probe complementary to the *qseB* region. Graph depicts log2(fold change) of *qseB* transcripts at each time point relative to the parental VUTI163 (mean ± SEM, n = 3). **(F)** VUTI247 and passage 2 and 5 were tested for activity of the *qse* operon. Cells were collected from a culture grown overnight allowed to reach stationary phase. RNA was extracted and reverse –transcribed. cDNA resulting from the reaction was subjected to qPCR with a probe complementary to the *qseB* region. Graph depicts log2(fold change) of *qseB* transcripts at each time point relative to the parental VUTI163 (mean ± SEM, n = 1).

We then asked whether we can identify clinical isolates re-wired to harbor subpopulations with induced *qseB* expression, or increased resistance to positively charged antibiotics. A total of 400 clinical urinary isolates from the Vanderbilt University Clinical Microbiology were tested using the standard broth microdilution method (**Figure 7B**) and an E-test strip method in which a gradient concentration of antibiotic is secreted into the agar medium (**Figure 7C**). Of these isolates, 4 exhibited hetero-resistance to PMB; the isolate consisted of a majority population with an MIC less than 1.0mg/L, and a minority population with significant resistance to PMB (**Figure 7B**). Isolation and subsequent sequencing of the resistant subpopulations from 2 out of the 4 isolates determined mutations in *sucAB*, glutamate racemase (*murI*) and *gadX* (**Figure 7B**), which codes for the regulator of the *gadAB* genes (**Figure 4**). In parallel studies, *in vitro* evolution of the susceptible subpopulation from each isolate in 0.75X the PMB MIC, led to increased PMB resistance by passage 2 (**Figure 7C-D**). Subsequent sequencing of passage 2- and passage 5 “resistant daughters” revealed the acquisition of mutations in the promoter region of *gadX* for both VUTI163 and VUTI247 (**Figure 7E** and **Supplementary File 2**). Transcript evaluation of these strains also indicated elevated expression of the *qseB* locus (**Figure 7G-H**). Together, these observations indicate that permanent rewiring in QseB-regulated anaplerotic pathways are identified in clinical isolates and may contribute to treatment failure in the clinic.

## Discussion

Antibiotic resistance among bacterial pathogens has become a global health threat, fueling the efforts to develop new antimicrobial treatments. Bacteria can mount resistance to antibiotics through acquisition of mobile genetic elements, namely through acquisition of plasmids encoding antibiotic resistance cassettes. In many pathogens, resistance to antimicrobial agents is encoded chromosomally. In *Salmonella sp*. and *Escherichia coli* resistance to a last resort antibiotic, polymyxin B, is intrinsically encoded. In these systems, resistance to polymyxin B is mediated through two-component signal transduction systems that lead to the upregulation of lipid A modifiers. This process, while increasing the net charge of bacterial membrane and repelling cationic moieties, comes at a metabolic cost associated with diverting central metabolites to synthesize modified LPS.

Here we demonstrate a metabolic circuit under the control of the QseB transcription factor that controls oxoglutarate homeostasis during the cellular response to positively charged antibiotics. Specifically, QseB controls the metabolism of by-products following amino-arabinose addition to nascent phospholipids by Arn* proteins which are activated by PmrA. ArnB is a direct target of both QseB and PmrA and catalyzes the conversion of undecaprnyl-4-keto pyranose to undecaprenyl 4-amin-4-deoxy-L-arabinose, a key step in the addition of an amino-arabinose moiety addition to a phospholipid. In this key catalytic step oxoglutarate is utilized, producing glutamate in the process. Glutamate is a critical commodity in the cell, being connected to glutamine and nitrogen metabolism (*36-40*). It is thus not surprising that the abundance of glutamate is under multi-level control, one that is monitored – as we now show – by QseB.

QseB directly targets and controls several genes involved in the anaplerosis of oxoglutarate. The importance oxoglutarate in sustaining modified LPS biogenesis is emphasized by the observations that exogenous oxoglutarate rescues the *qseB* deletion mutant during antibiotic treatment. This not only underscores the need for oxoglutarate during LPS modification, but also implicates the need for QseB’s metabolic control of oxoglutarate production. Metabolomic analyses tracking the abundance of glutamate, aspartate and Co-enzyme A (**Figure 5**) – combined with the transcriptional profiling results (**Figure 3**), point towards the use of glutamate through the pantothenate pathway and co-enzyme A production, which can then enter the TCA cycle either as acetyl-CoA or succinyl-CoA (**Figure 4**). Likewise, conversion of glutamate to fumarate via the *arg* gene products would supply fumarate. Intriguingly, our data point towards the conversion of glutamate to GABA through the function of the *gab/gad* gene products, for eventual conversion into succinate and re-introduction into the TCA cycle. This step would replenish succinate, bypassing the need to convert oxoglutarate to succinate via the *sucAB-* and *sucDC-* encoded complexes (**Figure 4**). Typically, GABA produced via the function of GadA/GadB and is exported from the cell via the action of the GadC exporter. The gene organization of *gadA* and *gadBC* is such that *gadBC* are transcribed from the same promoter, but transcripts are further processed at the *gadBC* intergenic region to release *gadB* and *gadC* transcripts (*41*). In our transcriptional analyses, *gadA* and *gadB* were among the most highly induced genes, but *gadC* transcript was not significantly altered (**Figure 3** and **supplementary file 1**). This indicates the presence of an additional mechanism that prevents GABA export so that it could potentially be used to produce succinate (**Figure 4**). The discovery of clinical isolates that have resistant subpopulations with mutations in *gadX*, along with the *in vitro* evolution results lends further support that this pathway can be exploited to confer resistance to polymyxin. Notably, none of the source patients were treated with polymyxins prior to isolation of the strains, as polymyxins are a last-resort antibiotic not deployed to treat UTI in the US. The presence of resistant subpopulations with the ability to resist polymyxins, combined with the *in vitro* evolution experiments, suggests that different envelope stresses may lead to mutations that alter the LPS, thereby collaterally producing antibiotic-resistant bacteria. Resistance to polymyxin can be obtained quickly, and also exploits the QseB-mediated metabolic pathway. We are in the process of quantifying GABA, succinate and fumarate in response to antibiotic stress in wild-type pathogenic *E. coli* and isogenic *qseB*-deletion and complemented strains and will be investigating the fitness costs and benefits of the resistant subpopulations during infection.

We found that QseBC-PmrAB signaling is prevalent across different strains of tested pathogenic *E. coli* including those that inhabit both intestinal and extraintestinal niches. This suggests that the mechanisms studied in this paper may be applicable across the major clades of *E. coli*. Reports of plasmids *mcr*-1 thru *mcr*-8 report low levels of polymyxin B and colistin resistance (*42, 43*). These elements lack *qseBC*. However, a new plasmid, *mcr*-9, encoding both a *mcr-*1 homologue and *qseBC* like elements suggests the need for metabolic control during LPS modification (*19*). In studying emerging antibiotic resistance mechanisms, we tend to generally focus on understanding plasmid-encoding systems. Here, understanding a chromosomally encoded system may translate to emerging plasmid encoded systems, and pose a threat in the clinic. Currently there are two polymyxins used clinically: polymyxin B and E (also known as colistin). Though highly effective, polymyxins fell out of favor in clinical practice due to concerns of renal and neurotoxicity in 1970s (*44*). Through the 1980s and 1990s, polymyxin use was largely restricted to cystic fibrosis populations (*45, 46*). However, the use of polymyxins in the clinic was reimplemented with the rise of multi-drug resistant infections in the early 2000s, where its use continues (*47, 48*). Likewise, aminoglycoside antibiotics such as gentamicin and amikacin are used to treat serious infections caused by facultative anaerobes (*49*). The finding that *E. coli* and potentially other Enterobacteriaceae have the potential to mount a resistance response – at the subpopulation level – to these critical antibiotics raises the alarm for the need to better understand mechanisms that lead to heterogeneous induction of systems like QseBC in the bacterial pathogens. Lastly, this work demonstrates the need to understand how metabolic pathways can be exploited in pathogenic bacteria and may give new insights to potential therapeutic targets.

## Materials and Methods

### Bacterial Strains, Plasmids, and Growth Conditions

Bacterial strains, plasmids and primers used in this study are listed in the **Table 2**. Overnight growth was always performed in liquid culture in Lysogeny Broth (Fisher) at 37°C with shaking, with appropriate antibiotics where necessary. Details pertaining to growth conditions for each assay used in the study can be found in the relevant sections below.

#### Transcriptional Surge Experiments

To assess induction of *qseBC*, bacteria were grown in N-minimal media (*27, 28*) at 37°C with shaking (220 rotations per minute). N-minimal media were inoculated with strains of interest at starting optical density at a wavelength of 600 nm (OD_600_) of 0.05. Strains were allowed to reach mid-logarithmic growth phase (OD_600_ = 0.5). At this time, 4-milliliters of culture was withdrawn for processing (see below) and the remainder of the culture was split into two. To one of the two split cultures, ferric chloride (Fisher) was added at a final concentration of 100 μM, while the other culture served as the unstimulated control. Cultures were returned to 37°C with shaking. Four-milliliter samples were withdrawn from each culture at 15- and 60- minutes post stimulation for RNA processing as described below. All samples were centrifuged at 4000 x g for 10 minutes upon collection. The supernatant was decanted and the fraction containing the cell pellet was flash frozen in dry ice – ethanol and stored at -80°C until RNA extraction.

### RNA Isolation

RNA from cell pellets was extracted using the RNeasy kit (Sigma Aldrich) and quantified using Agilent Technology (Agilent). A total of 3 micrograms (μg) of RNA was DNAse treated using the DNAfree kit (Ambion) as we previously described (*16, 20, 26*). A total of 1 μg of DNAse-treated RNA was subjected to reverse-transcription using SuperScript III Reverse Transcriptase (Invitrogen/ThermoFisher Scientific) and following the manufacturer’s protocol.

### RNA Sequencing and Analysis

Strains were grown in N-Minimal media at 37 °C with shaking, and samples were obtained as described for the transcriptional surge experiments. RNA was extracted and DNAse-treated as described in the RNA isolation section. DNA-free RNA quality and abundance were analyzed using a Qubit fluorimeter and Agilent Bioanalyzer. RNA with an integrity score higher than 7 was utilized for library preparation at the Vanderbilt Technologies for Advance Genomics (VANTAGE) core. Specifically, mRNA enrichment was achieved using the Ribo-Zero Kit (Illumina) and libraries were constructed using the Illumina Tru-seq stranded mRNA sample prep kit. Sequencing was performed at Single Read 50 HT bp on an Illumina Hi Seq2500. Samples from three biological repeats were treated and analyzed. Gene expression changes in a given strain as a function of time (15 minutes post stimulation versus unstimulated; 60 minutes post stimulation versus unstimulated) were determined using Rockhopper software hosted on PATRIC database.

### chIP-on-chip

To determine promoters bound by QseB, strain UTI897*qseB* was used, that harbors a construct expressing Myc-His-tagged QseB under an arabinose-inducible promoter (*12, 15, 27*). As a control for non-specific pull-downs, an isogenic strain harboring the pBAD-MycHis A empty vector was used. Cultures were grown in Lysogeny Broth in the presence of 0.027M arabinose to ensure constant expression of QseB, at concentrations similar to those we previously published as sufficient for QseBC complementation (*15*). Formaldehyde was added to 1% final concentration, following the methodology as described by Mooney et al., (*50*). Upon addition of formaldehyde, shaking was continued for 5 min before quenching with glycine. Cells were harvested, washed with PBS, and stored at -80 °C prior to analyses. Cells were sonicated and digested with nuclease and RNase A before immunoprecipitation. Immunoprecipitation was performed using an anti-Myc antibody (ThermoFisher) on six separate reactions, three for the experimental and three for the control strain. The ChIP DNA sample was amplified by ligation-mediated PCR to yield >4 μg of DNA, pooled with two other independent samples, labeled with Cy3 and Cy5 fluorescent dyes (one for the ChIP sample and one for a control input sample) and hybridized to UTI89-specific Affymetrix chips (*26*).

#### Antibiotic Survival Assays

To assess susceptibility of strains to polymyxins and aminoglycosides, strains were grown in N-minimal media in the absence (unstimulated) and presence (stimulated) of ferric iron (at a final concentration of 100μM) as described for the transcriptional surge experiments and in Figure 2. When bacteria reached an OD_600_ of 0.5, they were normalized to an OD_600_ of 0.5 in 5ml of 1X phosphate buffered saline (PBS)and split into two groups: A) Nothing added B) antibiotic added at a final concentration of 2.5 μg/mL for polymyxin B (5X MIC), 3.125 μg/mL for gentamicin (1X MIC), and 10 μg/mL for amikacin (5X MIC). The stimulated samples also received ferric iron at a final concentration 100μM. Samples were incubated for 60 or 180 minutes at 37 °C during which samples at 0, 15, 60 and 180 minutes post antibiotic addition or at only 60 minutes post antibiotic addition were serially diluted and plated on nutrient agar plates (Lysogeny Broth agar) to determine colony forming units per milliliter (CFU/mL). Percent survival as a function of ferric iron pre-stimulation for samples incubated for one time point, 60 minutes was determined by dividing the treated samples by CFUs of the unstimulated control sample and multiplying by 100. Percent survival as a function of ferric iron pre-stimulation for samples incubated for 180 minutes in which multiple samples at 0, 15, 60 and 180 minutes was determined by dividing the treated flasks by CFUs of the T = 0 sample of each condition and multiplying by 100. For polymyxin B (PMB) survival assays performed concurrently with metabolite measurements (see relevant section below), samples were taken across time (at induction (t=0), 15, 60, and 180 minutes post ferric iron additions). For PMB survival assays performed concurrently with oxoglutarate rescue assays, oxoglutarate (5 mM) was added at the same time as polymyxin B or gentamicin after standardization of the samples to an OD_600_ = 0.5

### Metabolite Measurements

Pellets of approximately 10^8^ cells were collected from PMB survival assays at each time-point. Glutamate, aspartate and coenzyme A levels were quantified using a colorimetric assay utilized from Glutamate-, Aspartate- and Coenzyme A Assay Kits (all kits obtained from Sigma Aldrich) utilizing the entire, undiluted sample (10^8^ cells). Assays were performed according to manufacturer’s instructions in at least 2 biological replicates per strain, per timepoint.

### Minimum Inhibitory Concentration (MIC) Determination

To determine the minimum inhibitory concentration of PMB, amikacin, and gentamcin in strains used in this study the broth microdilution method was used. Strains were grown at 37°C overnight with shaking, in cation-adjusted Mueller Hinton broth following clinical microbiology laboratory standard operating procedures (*51*). Specifically, strains were sub-cultured at a starting OD_600_ of 0.5 and allowed to reach growth at an OD_600_ = 0.4 – 0.5. Cells were then normalized to and OD_600_ = 0.5 or roughly 10^5^ cells. At this time, a 96 well polypropylene plate (for PMB) or a polystyrene plate (amikacin and gentamicin) was prepared with a gradient of concentrations (2-fold dilution) across the rows, plus a column with no antibiadded as a growth control, and a media only column to serve as a negative control. Five microliters of the standardized culture were added to each well except those holding the media control. Plates were incubated statically at 37°C for 24 hours. At this time, the minimum inhibitory concertation was determined. The concentration of antibiotic of the well in which bacterial growth was diminished by greater than 90% was determined to be the minimum inhibitory concentration if all three technical replicates were in agreement. Each strain was tested with 3 technical replicates and 3 biological replicates.

### Isolation and whole-genome sequencing of heteroresistant subpopulations from clinical isolates

*E. coli* was isolated from patient urine samples and banked by MicroVU in 25% glycerol freezer stocks. Strains were grown at 37°C overnight with shaking in lysogeny broth (Fisher). Strains were then diluted to OD_600_ = 0.5 in PBS, and adjusted to a 0.5 McFarland standard. Samples were then streaked out on prepared Mueller Hinton agar plates (BD) to create a lawn, using a cotton applicator (Puritan). A single polymyxin E-test strip (Biomerieux) was placed onto the lawn. Plate were incubated at 37°C for 16 hours. From samples in which two populations were phenotypically present on the plate after incubation: VUTI149, 163, 184 and 247, each population (sensitive and resistant) was isolated and grown overnight at 37°C in lysogeny broth. The samples were then prepared for genomic DNA isolation (Qiagen). Isolated genomic DNA from both populations were sequenced (GeneWiz). Briefly, libraries were prepared using the NEBNext UltraT DNA Library Prep Kit for Illumina. The genomic DNA was fragmented by acoustic shearing with a Covaris LE220 instrument. Fragmented DNA was end repaired and adenylated. Adapters were ligated after adenylation of the 3’ ends followed by enrichment by limited cycle PCR. DNA library was validated using a D100 ScreenTape on the Agilent 4200 TapeStation and was quantified using Qubit 2.0 Fluorometer. The DNA library was also quantified by real time PCR, clustered on a flow-cell and loaded on the Illumina Miseq instrument. Resulting reads were assembled using the Genome Assembly tool using SPAdes hosted on the Patric Databse. Genomes were then annotated using the *Escherichia* group as reference using the Annotation tool hosted by Patric. To determine variation between both subpopulations (sensitive and resistant), the reads of the resistant subpopulation were compared to the assembled genome of the sensitive population using the Variation Analysis tool hosted on Patric’s database.

### In vitro evolution of clinical isolate with polymyxin

For clinical isolates VUTI 163 and 247 an *in vitro* evolution experiment was performed. Strains were grown overnight at 37°C shaking. The next day, strains were sub-cultured 1:1000 in fresh Mueller Hinton broth with the addition of a sublethal amount of polymyxin, at 0.75X the MIC. Samples were sub-cultured 1:1000 in fresh Muller Hinton broth and polymyxin every 24 hours for 5 total passages. Genomic DNA from the second and fifth passage of VUTI63 and 247 were isolated (Qiagen) and sequenced (GeneWiz). Briefly, libraries were prepared using the NEBNext UltraT DNA Library Prep Kit for Illumina. The genomic DNA was fragmented by acoustic shearing with a Covaris LE220 instrument. Fragmented DNA was end repaired and adenylated. Adapters were ligated after adenylation of the 3’ ends followed by enrichment by limited cycle PCR. DNA library was validated using a D100 ScreenTape on the Agilent 4200 TapeStatio and was quantified using Qubit 2.0 Fluorometer. The DNA library was also quantified by real time PCR, clustered on a flowcell and loaded on the Illumina Miseq instrument. Resulting reads were assembled using the Genome Assembly tool using SPAdes hosted on the Patric Databse. Genomes were then annotated using the *Escherichia* group as reference using the Annotation tool hosted by Patric. To determine variation between the passages and parental strain, the reads of the resistant subpopulation were compared to the assembled genome of the sensitive parental population using the Variation Analysis tool hosted on Patric’s database.

### Quantification and Statistical Analysis

For polymyxin B survival assays, the percent survival of strains in specific conditions were calculated (mean ± SEM, N =3) and were compared to a control strain using Welch’s T-test of sensitivity performed using Prism software. For transcriptional surge experiments across time, no statistical test was used, but the mean ± SEM was displayed. For metabolite measurements, no statistical test was used, a representative of three biological replicates was displayed. For polymyxin B survival assays across time, no statistical test was used. For analysis of minimum inhibitory concentrations across passages in clinical isolates no statistical test was used, but the mean ± SEM was displayed.

## Supporting information

Supplementary File 1 - RNAseq data

Supplementary File 2 - WGS data

## Data and Code Availability

For whole genomes sequenced during this experiment, the sequencing data can be found on the Sequence Read Archive at PRJNA625502. RNA sequencing data submission can be found on ArrayExpress at E-MTAB-9277. ChIP-on-chip data can be found in supplementary file S2 and is pending submission at ArrayExpress.

## Author Contributions

MNH, CJB and MH designed, executed and interpreted experiments, prepared figures and wrote the manuscript. TS performed statistical analyses of RNAseq output and edited the manuscript. KRG designed and performed experiments and edited the manuscript. KAF created the *kguRS* deletion mutant. TB, AH, RM, SAR and DW performed and analyzed experiments and edited the manuscript.

## Acknowledgements

The authors would like to acknowledge the Center for Innovative Technologies for providing metabolomics support; the laboratory of Dr. Scott J. Hultgren for supporting the chIP-on-chip experiments through the following sources of funding: P50 DK64540, R01 AI048689 and R01AI02549; Dr. Erin J. Breland for helpful discussions regarding the transcriptional profiling; and the following sources of funding: R01 AI 5R01AI107052 and 1P20DK123967-01 to MH. Melanie Hurst is supported by NRSA F31 fellowship <u>1F31AI143244</u>-01A1.

