## Supplementary Figures S1-S4 for "A Bacterial Signaling Network Controls Antibiotic Resistance by Regulating Anaplerosis of 2-oxoglutarate"

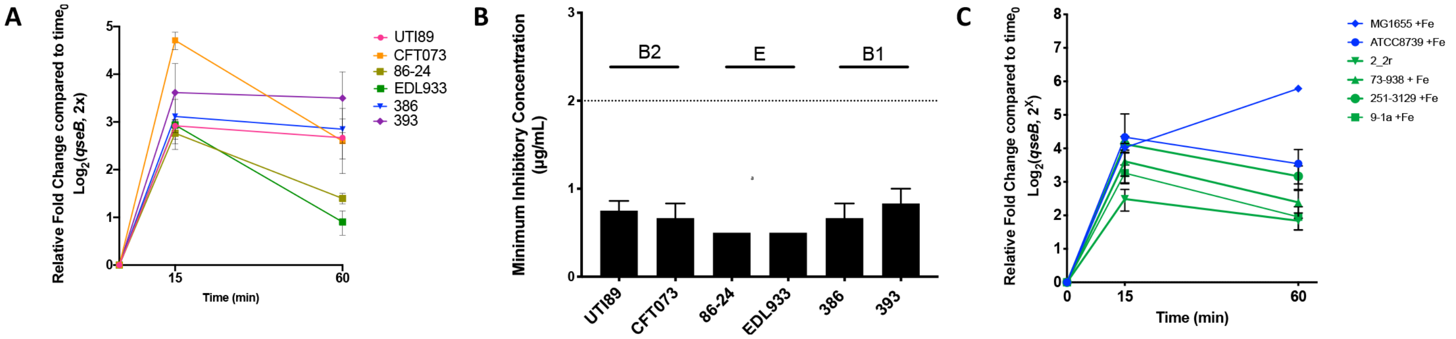


**Figure S1:** (A) Graph depicts the polymyxin B minimum inhibitory concentration (MIC) determined for the strains of pathogenic *E. coli* used in our studies. Strains from the most prevalent phylogenetic clades (depicted above the MIC of each graph by horizontal lines) are used (mean ± SEM, n = 3). (B) Graph depicts qPCR results tracking the activation of the *qse* operon following activation with ferric iron. Briefly, cells were allowed to reach mid-log growth phase. Cells were then collected before and at 15 and 60 minutes post addition of ferric iron. RNA was extracted and reverse –transcribed. cDNA resulting from the reaction was subjected to qPCR with a probe complementary to the *qseB* region. Graph depicts $\log_2$-fold change of *qseB* transcripts at each time point relative to the sample taken before stimulation (mean ± SEM, n = 3). (C) Strains from *E. coli* phylogenetic clades A (blue lines) and D (green lines) were tested for activation of the *qse* operon following activation with ferric iron. Cells were collected before stimulation and at 15- and 60-minutes post addition of ferric iron. RNA was extracted and reverse –transcribed. cDNA resulting from the reaction was subjected to qPCR with a probe complementary to the *qseB* region. Graph depicts $\log_2$(fold change) of *qseB* transcripts at each time point relative to the sample taken before stimulation (mean ± SEM, n = 3).

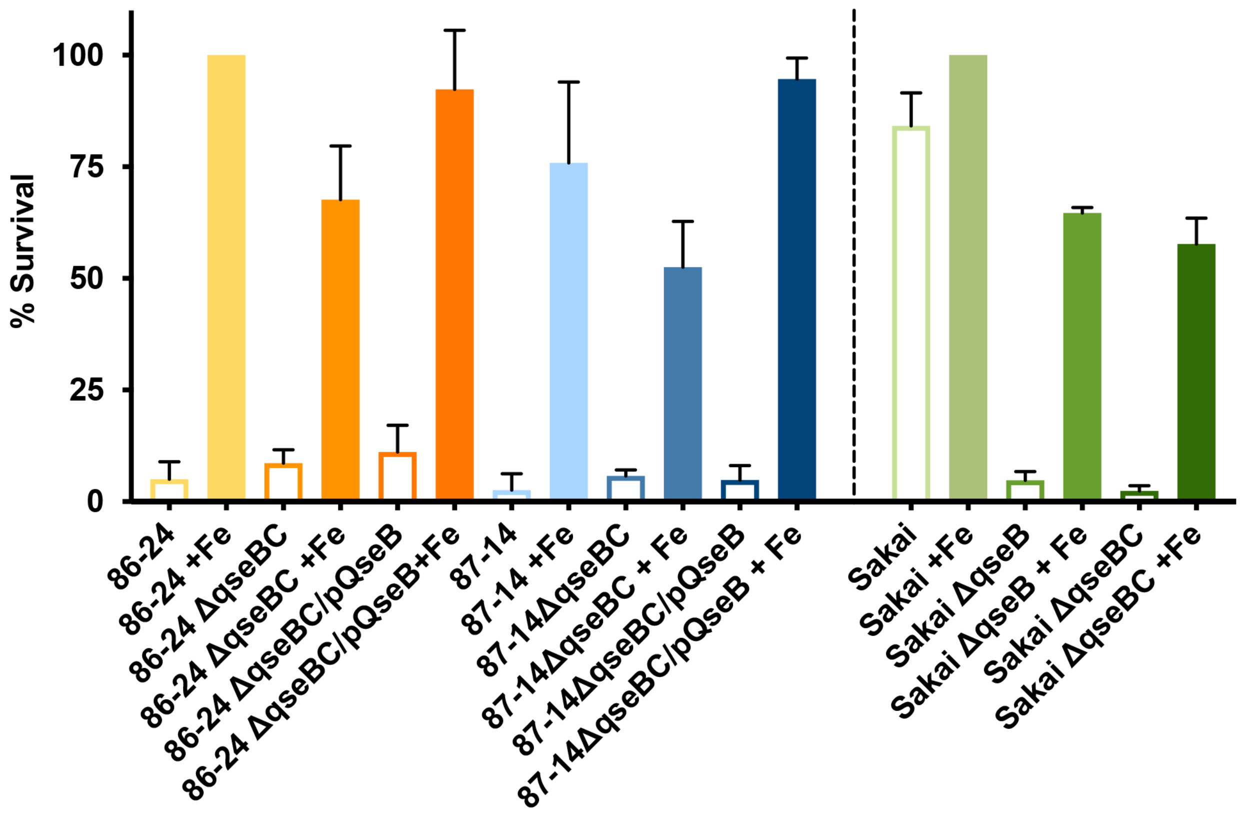


**Figure S2:** Graph depicts results of polymyxin B survival assays completed for representative EHEC strains and isogenic mutants. Cells were allowed to reach mid logarithmic growth phase in the presence or absence of ferric iron and normalized. Cells were then either exposed to polymyxin at 2.5 µg/mL or without addition for one hour. At this time cells were serially diluted and plated to determine colony forming units per mL. To determine percent survival, cells exposed to polymyxin were compared to those that were not (mean ± SEM, n = 3).

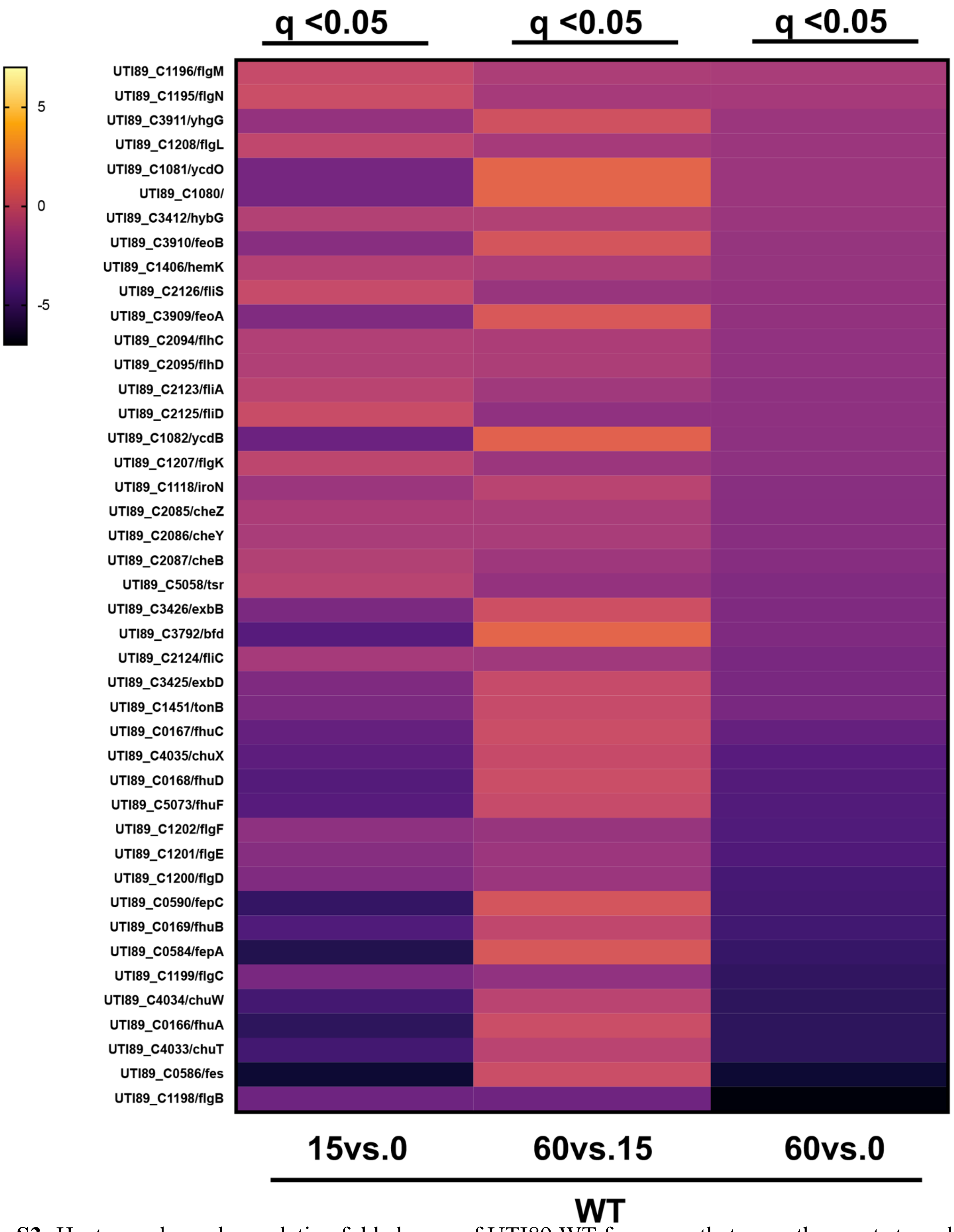


**Figure S3:** Heatmap shows $\log_2$ relative fold change of UTI89 WT for genes that were the most strongly down-regulated after stimulation with ferric iron at 15- and 60-minutes post stimulation. These genes were significantly ($q<0.05$) changed at 60 minutes compared to pre-stimulation (T=0) in wild-type cystitis isolate UTI89

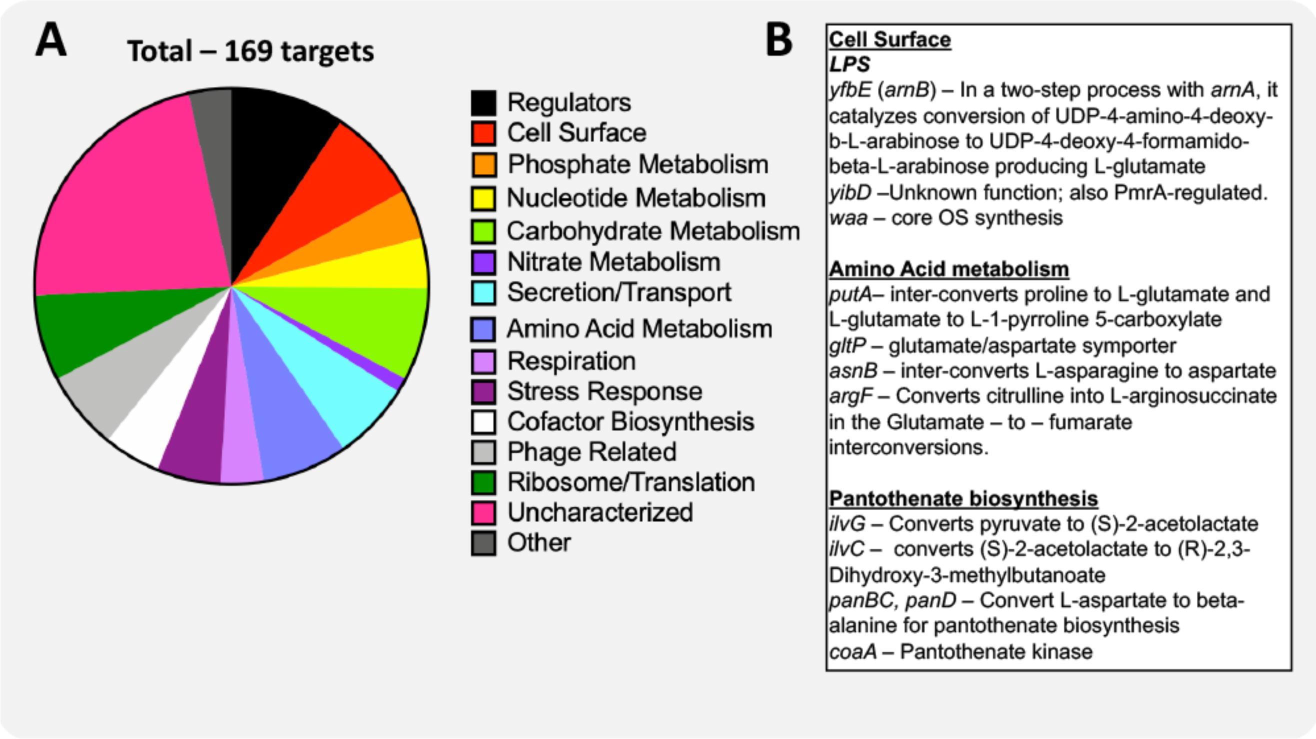


**Figure S4:** (A) Pie chart indicates the distribution of 169 unique DNA promoter sequences bound by QseB in pull-down experiments using tagged QseB and cross-linking, followed by immuno-precipitation, reversal of the crosslinks and hybridization of eluted DNA onto Affymetrix UTI89-specific chips. The data are from three independent biological experiments and exclude non- specific targets isolated through immunoprecipitation with vector control. (B) Subset of the direct targets of QseB that are differentially expressed following stimulation with ferric iron and are associated with glutamate metabolism.
